# Ablation of hepatocyte derived-FGL1 does not aggravate metabolic dysfunction-associated steatotic liver disease and hepatocellular carcinoma

**DOI:** 10.1101/2024.12.23.628701

**Authors:** Jean Personnaz, Lisa Cannizzo, Céline Marie Pauline Martin, Aurore Desquesnes, Manon Sotin, Joanna DaSilva, Prunelle Perrier, Hervé Guillou, Léon Kautz

## Abstract

Metabolic dysfunction-associated steatotic liver disease (MASLD) begins with simple steatosis, which can progress to hepatocellular carcinoma (HCC). The pathogenesis of MASLD alters the secretion of hepatokines such as fibrinogen-like 1 (FGL1), a candidate mediator of liver steatosis and hyperglycemia. To investigate the contribution of FGL1 to liver diseases, we compared wild-type mice to mice with hepatocyte-specific deletion of *Fgl1* subjected to a steatosis or HCC experimental protocol. We found that mice deficient for *Fgl1* in hepatocytes showed higher levels of plasma glucose, pronounced metabolic alterations and liver injury when fed a western diet compared to their wild-type counterparts. However, both genotypes exhibited a similar lipid deposition in the liver. Similarly, wild type and *Fgl1*-deficient mice displayed comparable liver alterations during HCC progression. We observed that *FGL1* expression was repressed during MASLD progression in mice and human concomitantly with the severity of liver injury. Altogether, these findings suggest that FGL1 is not a major contributor to the pathogenesis of MASLD and HCC.

## INTRODUCTION

Metabolic dysfunction-associated steatotic liver disease (MASLD), previously known as non-alcoholic fatty liver disease (NAFLD) (1), initiates from an asymptomatic steatosis affecting 30% of the worldwide population (2). Approximately 15% of steatotic patients develop a metabolic dysfunction-associated steatohepatitis (MASH, previously known as NASH), defined by the installation of fibro-inflammatory phenomenons, hepatocytes ballooning and steatosis. The MASH stage represents a critical turning point, as 24% of patients with MASH progress to cirrhosis within 8 years, and 3% are diagnosed with hepatocellular carcinoma (HCC) annually (3). MASLD currently represents the fastest growing cause of HCC worldwide (4) and a significant economic burden (2). MASLD is the hepatic manifestation of the metabolic syndrome characterized by obesity, insulin resistance, hypertension, increased lipolysis and dyslipidemia (5, 6). Excess of lipolysis-derived circulating fatty acids are then stored as triglycerides in the liver thereby causing steatosis (6, 7). Fat accumulation in hepatocytes triggers transcriptional modifications and secretion of hepatokines with various metabolic effects in fatty livers (8). Among those, previous studies suggested that the induction of Fibrinogen-like 1 (FGL1)/ hepassocin during MASLD contributed to disease progression, insulin resistance and hyperglycemia (9–11).

FGL1 is a 34kDa protein secreted exclusively by hepatocytes (12) that shares a structural homology with fibrinogen β and γ chains. FGL1 is not involved in the clotting process as it lacks the platelet binding domain, the cross-linking region and the thrombin sensitive site (13). In mice, the manipulation of FGL1 has yielded contradictory results. Administration of FGL1 after the induction of hepatic injury by 3 weeks of methionine and chlorine-deficient diet reduced hepatic triglycerides content and fibrosis (14). Conversely, mice fed and standard chow and treated with recombinant FGL1 were less sensitive to insulin (9). *Fgl1* silencing using RNA interference was protective against high fat diet induced liver injury (10) whereas the overexpression of *Fgl1* was sufficient to induce hepatic steatosis. However, *Fgl1-*deficient mice fed a standard rodent chow displayed an increased body weight and a mildly increased fasting hyperglycemia without any modification of food intake (14, 15) compared to wild type mice suggesting that FGL1 could be involved in the onset of obesity. Moreover, *Fgl1*-/- mice developed a more severe hepatocellular carcinoma induced by diethynitrosamine (DEN) (16). Interestingly, *FGL1* gene is located on chromosome 8 both in human and mouse in a region containing several tumor suppressor genes such as *VPS37A* or *DLC1* (17). FGL1 has also been described as a mediator of immune evasion in certain cancers (18) but this function has since been questioned (19–21). Overall, the contribution of FGL1 in the progression of MASLD remains elusive. Here, we compared the MASLD and HCC phenotype of WT mice (*FGL1*^LWT^) and mice with hepatocyte-restricted deletion of *Fgl1 (Fgl1*^LKO^). We observed that *Fgl1*^LKO^ mice exhibited increased circulating levels of liver transaminases and a trend toward a higher incidence of solid tumors compared to *FGL1*^LWT^ mice. However, we found no differences in steatosis or fibrosis. Furthermore, in contrast to published studies, we observed decreased FGL1 levels in both mice and humans during disease progression.

## MATERIAL AND METHODS

### Animals

#### Fgl1 liver knock out mice

Fgl1^tm1b(EUCOMM)Hmgu^ mice were obtained from the international mouse phenotyping consortium (22) and crossed with mice expressing the FLPO recombinase (C57BL/6-Tg(CAG-Flpo)1Afst/Ieg; MMRRC_036512-UCD) to remove the EUCOMM cassette flanked by FRT site resulting in *Fgl1* floxed (*Fgl1^fl/fl^)* mice. *Fgl1^fl/fl^* mice were then bred with mice expressing the CRE recombinase under the control of the albumin promoter (B6.Cg-Speer6-^ps1Tg(Alb-cre)21Mgn/J^, Jackson laboratory stain #003574) to obtain mice with hepatocytes specific ablation of *Fgl1 (Fg1*^LKO^) or wild type *Fgl1^fl/fl^* littermate mice *(Fg1l*^LWT^) (23). Mice were bred and housed at Janvier Laboratories (Le Genest St Isle, France) and transferred to the CREFRE US006 animal care facility at 4 to 5 weeks of age. 6 week-old C57BL/6J were purchased from Envigo. Mice were housed in a specific and opportunistic pathogen-free facility under a 12-hour light - dark cycle with *ad libitum* access to water and standard laboratory mouse chow diet (SAFE A04), in accordance with the European Union guidelines, and then subjected to a dietary challenge between 6 to 8 weeks of age. A set of *Fgl1*^LWT^ and *Fgl1*^LKO^ mice was fasted overnight (15h) prior to euthanasia. Liver and adipose tissue weight relative to total body weight was represented as liver or adipose tissue (%). The study was performed in compliance with the European guidelines for the use and care of laboratory animals, and approved by an independent ethics committee (CEEA 122) under the authorization numbers APAFIS #40791-202302021502788.

#### Steatosis model

*Fgl1*^LWT^ and *Fgl1*^LKO^ mice were fed a normal chow (SAFE A04) or a western diet containing 35% sucrose, 20% lipids and 0,15% cholesterol (Research diet D12079B) for 16 weeks with free access to water.

#### Oral glucose tolerance test

Mice were housed in clean cages without food for 15h. Baseline glycemia was first measured from a drop of blood harvested at the tail vein using a glucometer (FreeStyle Optium Neo, Abbott). 30 minutes later, mice were administered a 1.5g/kg bolus of a D-Glucose solution *per os.* Glycemia was recorded over 120 minutes.

#### Hepatocellular carcinoma model

*Fgl1*^LWT^ and *Fgl1*^LKO^ mice were fed a western diet containing 35% sucrose, 20% lipids and 0,15% cholesterol (Research diet D12079B) with weekly intraperitoneal injection of carbon tetrachloride (CCl_4_, 0.2µl/g) (24) for 24 weeks.

### Genotyping

After digestion with 25mM NaOH, 0.2M EDTA and neutralization with 40mM Tris-HCl, PCRs were performed using the following primers:

Alb-CRE:

F1-TGCAAACATCACATGCACAC,

F2-GAAGCAGAAGCTTAGGAAGATGG,

R-TTGGCCCCTTACCATAACTG,

LoxP sites:

F-TGTGCAGACTGTCATCTCAGTACAGCC,

R-TGTGCAGACTGTCATCTCAGTACAGCC.

The excision of *FGL1* in the hepatocytes was confirmed using primers flanking exon 2:

F-TGTGCAGACTGTCATCTCAGTACAGCC,

R-GGCAGAGCTCGAGCATTAGCAGA.

### Biochemical analysis

Whole blood was collected with K_3_-EDTA (Sigma) to prevent clotting and the plasma was prepared by two successive centrifugations (3000rpm at 4°C for 5min). Levels of triglycerides, cholesterol, alanine and aspartate aminotransferases were determined by the Anexplo Phenotyping facility using a Pentra400 biochemical analyzer (HORIBA Medical, Kyoto, Japan). Blood β-Hydroxybutyrate levels were measured from a drop of blood harvested at the tail vein using a FreeStyle Optium Neo (Abbott).

### Liver glycogen quantification

Frozen liver tissues were weighed, homogenized in 4% (v/v) ice-cold perchloric acid (50 mg/mL) and centrifuged for 10 min at 8,000 rpm and 4°C. Acid supernatant was neutralized and glycogen was hydrolyzed for 1 h at 55 °C with α-(1–4)-(1–6)-amyloglucosidase (10 mg/mL, Roche Applied Science). The amount of glucose released was quantified by measuring at 25 °C the appearance of NADPH at 340 nm: 250 µL final volume containing 100 mM Tris (pH 8.5), 15 mM MgCl2, 0.5 mM NADP, and 0.7 mM ATP was added, followed by the addition of 1 unit of glucose-6-phosphate dehydrogenase (Roche Diagnostics) for 10 min. After a first lecture at 340 nm, 1.5 units of hexokinase (Roche Diagnostics) were added for 15 min before measuring the final glucose released at 340 nm. Results were expressed as µmoles of glycogen per µg of proteins per milligram of liver.

### Neutral lipids analysis

Lipids from 1mg of liver weight were extracted using adapted Bligh and Dyer method (25) in dichloromethane/methanol (2% acetic acid) /water (2.5:2.5:2 v/v/v). Stigmasterol, cholesteryl heptadecanoate and glyceryl trinonadecanoate were used as internal standards. The solution was centrifuged at 2500 rpm for 6 min. The organic phase was collected and dried under nitrogen, then dissolved in 20 µL of ethyl acetate. 1μl of the lipid extract was analyzed by liquid-gas chromatography on a Trace 1600 (Thermo Electron) system using a RTX-5 Restek fused silica capillary column (5mx0.25mm ID, 0.50 μm film thickness) (26). The oven temperature was programmed from 200°C to 350°C at a rate of 5°C per minute, and the carrier gas used was hydrogen (0.5 bar). The injector and detector temperatures were set at 315°C and 345°C, respectively.

### RNA extraction, RT and qPCR

Total RNA from mouse tissues was extracted by Trizol (MRC) / Chloroform (Sigma) method. Complementary cDNA was synthetized using M-MLV Reverse transcriptase (Promega). Messenger RNA (mRNA) expression levels were assessed by quantitative polymerase chain reaction (RT-qPCR) using Takyon SYBR green (Eurogentec) and run on a LightCycler480 (Roche) apparatus. Sequences of the primers used were:

*Scd1*: F-TGGAGACGGGAGTCACAAGA, R-ACACCCCGATAGCAATATCCAG;

*Fasn*: F-TACAACCTCTCCCAGGTGTG, R-CCTCCCGTACACTCACTCGT;

*Acta2:* F-CCAGCCATCTTTCATTGGGATG, R-AGCATAGAGATCCTTCCTGATGT;

*Col1a1*: F-CGATGGATTCCCGTTCGAGT, R-GAGGCCTCGGTGGACATTAG;

*Cd45*: F-AGGTGTCCTCCTTGTCCTGT, R-TGTACACACCCACAGCACTC;

*Emr1*: F-AAGGACACGAGGTTGCTGACC, R-GCCAATCTGGAAAATGCCC;

*Fgl1*: F-CGATCTGATGGCAGTGAGAACT, R-TTTGTTACCCAGCCAGTATTCG;

*Il6*: F-CTCTGCAAGAGACTTCCATCCAGT, R-CGTGGTTGTCACCAGCATCA;

*Ly6g*: F-CTCCTGCAAGCAGACAGTGAT, R-GGGTAGTGATGGCTCAAGGTC;

*Cd3e*: F-GGACAGTGGCTACTACGTCTGCT, R-CACACAGTACTCACACACTCGAGC;

*Ddit3*: F-AGCCAGAATAACAGCCGGAAC, R-TTCTGCTTTCAGGTGTGGTGG;

*Socs3*: F-TTAAATGCCCTCTGTCCCAGG, R-TGTTTGGCTCCTTGTGTGCC

*Pck1:* F-GGGGTGTTTGTAGGAGCAGC, R-GGCCAGGTATTTGCCGAAGT

*G6pc:* F-CTGCTCACTTTCCCCACCAG, R-GAATCCAAGCGCGAAACCAAA

*Rpl4*: F-TGAAAAGCCCAGAAATCCAA, R-AGTCTTGGCGTAAGGGTTCA.

Transcript abundance was normalized to the reference gene *Rpl4* and represented as a difference between reference and target genes within each group of mice (−ΔCt) ± standard deviation (SD).

### RNA sequencing

Pair-end RNA sequencing was performed by BGI DNA company (China). Briefly, total liver mRNA was isolated from 8 weeks-old *Fgl1*^LWT^ and *Fgl1*^LKO^ (n=7 per genotype). After quality control, bulk mRNA was sequenced using BGI DNBSEQ platform. After filtering, clean reads (150 bases) were aligned to mouse genome (mm39) using HISAT. Bowtie2 software was used to align the clean reads to the reference genes. Identification of differentially expressed genes was performed using Dr Tom software. RNAseq data and experimental details are available in NCBI’s Gene Expression Omnibus and are accessible through GEO Series accession number GSE291163.

### Production of recombinant FGL1

The production of Fc-tagged mouse FGL1 was carried out as previously described (27). *Fgl1* cDNA was cloned into pFUSEN-hG2Fc vector (Invivogen). Recombinant protein was produced in Freestyle 293F cells (Invivogen). Recombinant protein was purified using Hitrap Protein A HP column on an ÄKTA pure chromatography system (GE healthcare) and eluted with 0.1M Glycine pH 3.5. After concentration, FGL1 recombinant protein was resuspended in 0.9% NaCl. Protein purity and concentration were determined using Coomassie Imperial Protein Stain and Pierce bicinchoninic acid protein assay (Thermo Fisher Scientific).

### Primary hepatocytes

Hepatocytes were isolated form 8-12 weeks old wild type male C56BL/6J mice using a two-step collagenase perfusion as previously described (28). 4 to 6 hours after plating, cell media was changed to Williams’E media (ThermoFisher), 1% penicillin/streptomycin without serum supplemented with 10µg/ml of Fc fragment or recombinant FGL1 for 15h.

### Western Blot

Cell samples were lysed in RIPA buffer (25 mM Tris-HCl pH 7.6, 150 mM NaCl, 1% NP-40, 1% sodium deoxycholate, 0.1% SDS, ThermoScientific), supplemented with anti-protease (Roche) and anti-phosphatase cocktail 2 (Sigma). Liver biopsies were solubilized in protein extraction buffer (150mM NaCl, 5mM Tris-HCl, 0.5M EDTA, 0.1% NP40), supplemented with anti-protease (Roche) and anti-phosphatase cocktail 2 (Sigma). Protein concentration was assayed by Pierce bicinchoninic acid protein assay (Thermo Fisher Scientific) and 5µg of protein was loaded on a 4%-15% polyacrylamide gel (BioRad) and transferred on a nitrocellulose membrane. After blocking with Tris-buffered saline containing 2.5% BSA and 0.15% Tween 20 (TBST-BSA), membranes were incubated with the following antibodies from Cell Signaling Technologies diluted 1/1000 in TBST-BSA: phospho-AKT (Thr308, Rabbit mAb #13038), AKT (Rabbit mAb #9272), phosphor-ERK1/2 (Thr202/Tyr204, Rabbit mAb#4376) and ERK1/2 (Rabbit mAb#4695). Membranes were then probed with an HRP-linked anti-rabbit secondary antibody (Cell signaling technology). Enzyme activity was developed using ECL prime reagent (GE Healthcare) on ChemiDoc XRS+ imaging system. Phosphorylation levels were normalized to the signal intensity for corresponding non-phosphorylated proteins using Biorad Image Lab software and global quantitation tool.

### Histology

Liver biopsies were fixed in 4% formaldehyde for 24 hours before paraffin-embedding. 5 µm sections of paraffin-embedded livers were stained with hematoxylin and eosin (H&E, Sigma), picrosirius red (Sirius red, Abcam) or F4/80 (Cell signaling) according to standard protocols. A NAFLD activity score (NAS) (29) was computed.

### Statistical analysis

All data are expressed as mean ± standard deviation (SD). Statistical significance was assessed by Student t test, 1- or 2-way analysis of variance followed by post-hoc test as specified in the Figure legends using GraphPad Prism version 10. P values <0.05 were considered significant (*P < 0.05; **P < 0.01; ***P < 0.001; ****P < 0.0001).

## RESULTS

### Hepatocyte-specific deletion of *Fgl1* promotes body weight gain

While previous studies suggested that adipose tissue is a secondary source of FGL1, *Fgl1* expression was 200-fold lower in inguinal (iWAT) or epidydimal (eWAT) white adipose tissue than in the liver indicating that hepatocyte-derived *Fgl1* is the main source of plasma FGL1 (Figure 1A). *Fgl1* floxed mice were bred with mice expressing the CRE recombinase under the control of the albumin promoter to invalidate *Fgl1* specifically in hepatocytes (*Fgl1*^LKO^). Excision of *Fgl1* exon 2 in hepatocytes led to a 170-fold reduction in whole liver *Fgl1* mRNA expression compared to *Fgl1*^LWT^ mice (Figure 1B). We next compared the transcriptomic profiles of livers from *Fgl1*^LWT^ *and Fgl1*^LKO^ mice by RNA sequencing. We found that only 8 unrelated transcripts were differentially expressed between the two genotypes suggesting that FGL1 had virtually no effect on liver transcription (Figure 1C). Consistent with the phenotype of mice with constitutive deletion of *Fgl1* (15), 8 weeks old *Fgl1*^LKO^ mice displayed increased body weight and inguinal fat (iWAT) deposit compared to wild-type littermates (Figure 1D) but no difference in liver (Figure 1E) and epididymal (eWAT, Figure 1F) relative weight was observed.

**Figure 1:**
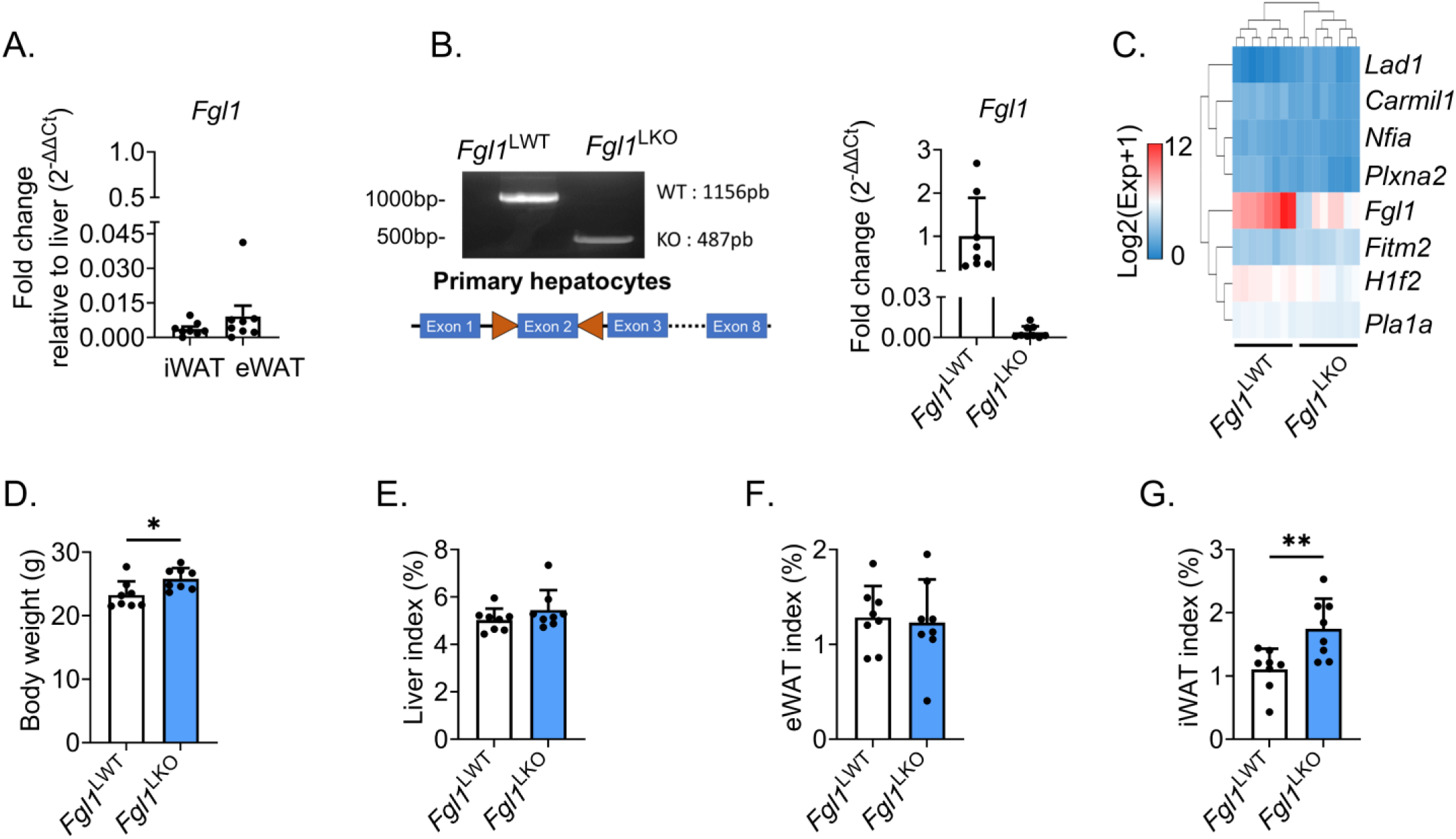
Hepatocyte specific deletion of *Fgl1* recapitulates the phenotype of whole-body knockout mice. Fold change in *Fgl1* mRNA expression inguinal adipose tissue (iWAT) and epididymal adipose tissue (eWAT) relative to the liver (A). Validation of *Fgl1* exon 2 excision in primary hepatocytes by PCR and qPCR (B). Heatmap of differentially expressed genes in the liver of *Fgl1*^LWT^ (n=8) and *Fgl1*^LKO^ (n=8) mice analyzed by mRNA sequencing (C). Body weight (D), liver (E), epididymal adipose tissue (eWAT, F) and inguinal adipose tissue (iWAT, G) weight relative to total body weight index (%) in *Fgl1^LWT^* (n=8) and *Fgl1*^LKO^ (n=8) males mice fed a normal chow. Data are mean ± SD. Statistical significance was assayed by Student’s t-test (B-G) or Two-way ANOVA, followed by Šídák’s multiple comparisons test (H). *p<0.05, **p<0.01, ***p<0.001

### Deletion of hepatocytes *Fgl1* does not significantly alter glucose metabolism

*Fgl1* whole-body knockout mice presented with elevated glucose circulating levels after starvation (14, 16). To determine whether hepatocyte FGL1 regulates glucose metabolism, *Fgl1*^LWT^ *and Fgl1*^LKO^ mice were subjected to a 15-hour fasting. The level of the ketone body β-hydroxybutyrate was comparable between genotypes after fasting (Figure 2A) but *Fgl1*^LKO^ mice exhibited a slightly increased glycemia and a trend toward a lower liver glycogen content (Figure 2B-C). The hepatic expression of phosphoenolpyruvate carboxykinase (*Pck1*) was similar between genotypes and we observed a modest yet-significant reduction in glucose-6-phosphatase (*G6pc*, Figure 2D) expression in the liver of *Fgl1*^LKO^ mice compared to *Fgl1*^LWT^ mice suggesting that the gluconeogenic activity was not altered by *Fgl1* ablation. The phosphorylation level of the kinases AKT and ERK1/2 that are activated by the insulin receptor was also comparable in the liver of *Fgl1*^LWT^ and *Fgl1*^LKO^ mice (Figure 2E-F). While *Fgl1^LKO^* animals exhibited an slightly increased fasting glycemia (Figure 2B), the glucose response was similar between genotypes during an oral glucose tolerance test (oGTT, Figure 2G). Collectively, these results indicate that the phenotype of mice with whole-body ablation of *Fgl1* (15, 30) is attributable to hepatocyte-derived FGL1. To examine whether FGL1 exerts autocrine effects on insulin signaling (9, 12, 31), primary mouse hepatocytes were treated for 15 hours with 10µg/ml of recombinant Fc fragment (hIgG2) or Fc-FGL1 in presence of insulin. FGL1 had no effect on the phosphorylation of ERK1/2 but a faster induction of AKT was observed (Figure 2H) suggesting that FGL1 could increase insulin sensitivity *in vitro*.

**Figure 2:**
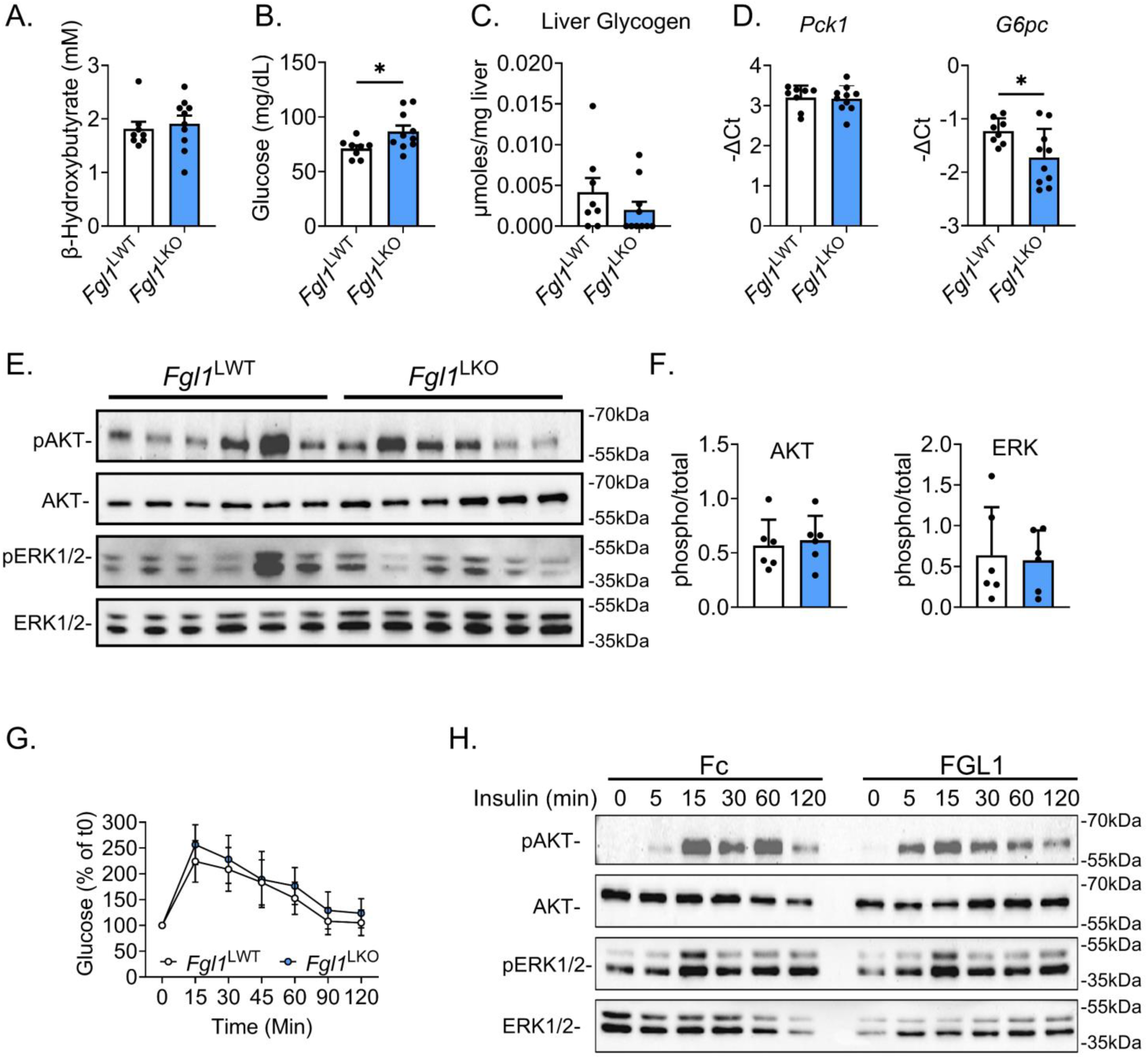
FGL1 does not have a major impact on glucose metabolism. Fasting blood β-hydroxybutyrate (A) and glucose (B) and liver glycogen (C) levels were measured in *Fgl1*^LWT^ or *Fgl1*^LKO^ mice (n=7 per genotype). *Pck1* and *G6pc* (D) mRNA expression (n=8) and (E) phosphorylation levels of AKT (Thr308) and ERK1/2 (Thr202/Tyr204) (n=6) in the liver of *Fgl1*^LWT^ and *Fgl1*^LKO^ mice. Total AKT and ERK were used as controls for quantification (F). Oral glucose tolerance test (1,5 g/kg, F) in *Fgl1*^LWT^ (n=6) or *Fgl1*^LKO^ (n=8) mice fed a normal chow (G). Phosphorylation levels of AKT (Thr 308) and ERK1/2 (Thr202/Tyr204) in hepatocytes stimulated with 100nM of insulin after a 15 hour treatment with recombinant Fc or FGL1 (10µg/ml) (H). Data are means ± SD and were compared by Student’s t-test (A-F) or Two-way ANOVA followed by Holm-Šídák’s multiple comparisons test (G), *p<0.05, **p<0.01, ***p<0.001.

### Deletion of hepatocyte *Fgl1* promotes metabolic alterations induced by western diet

Previous studies suggested that FGL1 could be involved in the pathogenesis of MASLD (9, 10, 14, 15, 32) but it is unclear whether this contribution is protective or detrimental. In order to investigate the contribution of hepatocyte-derived FGL1 in MASLD, 6-8 week-old *Fgl1*^LWT^ and *Fgl1*^LKO^ male mice were fed a normal chow or a western diet for 16 weeks to induce an obesity coupled with reduced glucose tolerance, hepatic steatosis and inflammation (33). Both genotypes gained a comparable weight when fed either a normal chow (Figure 3A) or a western diet (Figure 3B). 22-24-week-old *Fgl1*^LWT^ and *Fgl1*^LKO^ fed a normal chow or a western diet showed a comparable glycemia (Figure 3C-D). Glucose tolerance was similar between genotypes when mice were fed a normal chow but glucose levels were slightly elevated in *Fgl1*^LKO^ mice fed a western diet and subjected to an oral glucose tolerance test. Circulating triglycerides were unaffected by the dietary challenge but *Fgl1*^LWT^ and *Fgl1*^LKO^ mice fed a western diet showed a significant increase in circulating cholesterol compared to mice fed a normal chow (Figure 3E-F). Liver to body weight ratio was marginally increased (2%) in *Fgl1*^LKO^ mice fed a western diet in comparison to mice on normal chow but the ratios were similar between genotypes (Figure 3G). Circulating levels of alanine (ALAT) and aspartate aminotransferases (ASAT) (Figure 3H) were increased in mice fed a western diet compared to mice fed a normal chow and to a greater extent in *Fgl1*^LKO^ mice suggesting a higher susceptibility to diet-induced liver injury. Total liver expression of *Fgl1* was significantly reduced in *Fgl1*^LKO^ mice fed a standard or a western diet (Figure 3I) compared to their wild-type counterparts. *Pck1* mRNA expression was reduced in *Fgl1* ^LWT^ fed a western diet compared to mice fed a normal chow and *Fgl1*^LKO^ mice fed a western diet showed slightly elevated *Pck1* levels compared to their WT counterparts. iWAT relative weight was significantly increased only in *Fgl1*^LKO^ mice fed a western diet in comparison to mice on normal chow (Figure 3J) whereas the eWAT relative weight was significantly decreased in *Fgl1*^LKO^ mice compared to *Fgl1*^LWT^ mice (Figure 3L). Expression of *Fgl1* in iWAT (Figure 3K) or eWAT (Figure 3M) was low and unchanged by the dietary challenge in both genotype indicating that the absence of hepatocyte *Fgl1* is not compensated by the adipose tissue. Expression of genes involved in lipolysis *Lipe* (encoding the hormone-sensitive lipase) and *Pnpla2* (encoding the adipose triglyceride lipase) in the iWAT (Figure 3K) or the eWAT (Figure 3M) was comparable between groups and genotypes, suggesting that the lipolytic capacities of the adipose tissue deposits were not affected by the ablation of *Fgl1* in the liver. Lipoprotein lipase (*Lpl*) expression was significantly decreased in the iWAT of *Fgl1^LWT^* mice fed a WD compared to mice fed a NC (Figure 3K) but its expression in the eWAT was similar among the different groups (Figure 3M). The expression of the fatty acid transporter *Cd36* in the iWAT (Figure 3K) or in the eWAT (Figure 3M) was also unchanged. These results indicate that the deletion of *Fgl1* has a limited effect on MASH-related liver injury and adipose tissue function at this time point. We next analyzed liver tissue sections stained with hematoxylin and eosin (H&E) or sirius-red to assess the extent of liver damage. We observed that both *Fgl1*^LWT^ and *Fgl1*^LKO^ mice fed a western diet developed a comparable microvesicular steatosis in comparison to mice fed a normal chow (Figure 4A) whereas no sign of fibrosis was observed (Figure 4B). While the steatosis was comparable between *Fgl1*^LWT^ and *Fgl1*^LKO^ fed a western diet, a significant increase in non-alcoholic fatty liver disease activity score (NAS) was observed only in *Fgl1*^LKO^ mice compared to their controls (Figure 4C). We observed a significant and similar increase in hepatic cholesterol and triglycerides levels when mice were fed a WD (Figure 4D-E). A mild induction in lipogenesis marker *Scd1* was observed for both genotypes in mice fed a western diet compared to mice fed a normal chow while the expression of the inflammation and steatosis marker *Fasn* was unchanged (Figure 4F). Consistent with the absence of notable fibrosis, hepatic mRNA expression of *Col1a1* encoding the pro-α1 chain of procollagen or *Acta2* encoding the smooth-muscle actin was similar between animals fed a normal chow or a western diet (Figure 4G). mRNA expression of the inflammatory markers *Cd45* and *Emr1* (also known as F4/80, Figure 4H) was significantly increased in mice fed a WD but no difference was observed between *Fgl1*^LWT^ and *Fgl1*^LKO^. Altogether, these results suggest that the deletion of *Fgl1* has no to limited effect on MASH-related liver injury at this time point.

**Figure 3:**
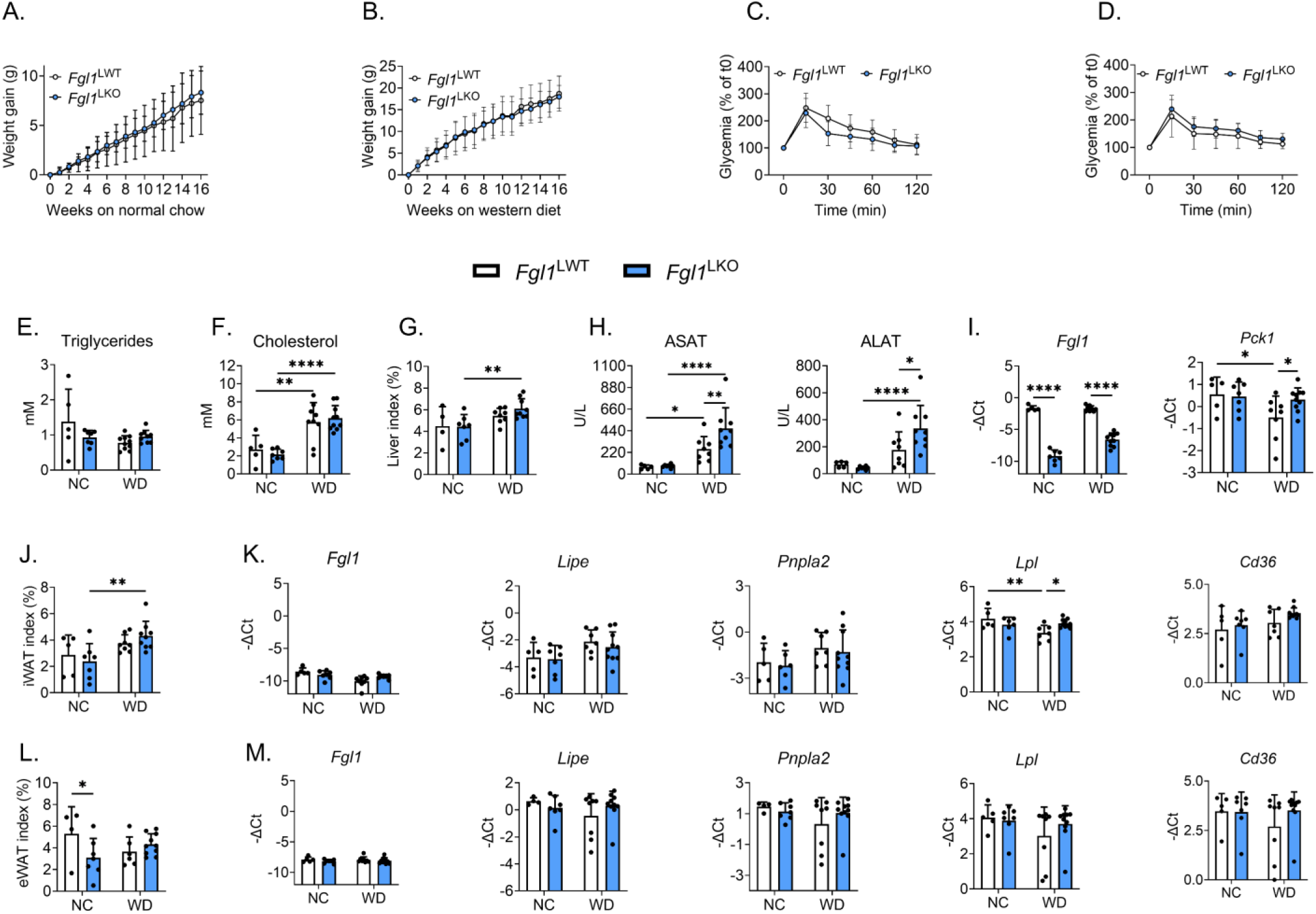
Deletion of *Fgl1* in hepatocytes sensitizes mice to diet-induced liver injury. *Fgl1*^LWT^ and *Fgl1*^LKO^ male mice were fed a normal chow or a western diet for 16-weeks. Body weight gain in mice fed a normal chow (A) or a western diet (B). Oral glucose tolerance test when mice were fed a normal chow (C) or a western diet (D). Circulating levels of triglycerides (E), cholesterol (F), liver weight relative to total body weight index (%) (G), aspartate (AST) and alanine (ALT) (H) aminotransferases, liver *Fgl1* and *Pck1* mRNA expression (I). iWAT (J) eWAT (L) weight relative to total body weight index (%) and mRNA expression of *Fgl1*, *Lipe, Pnpla2, Lpl* and *Cd36* (K-M). Data are means ± SD (*Fgl1^LWT^* NC n=5, *Fgl1^LKO^* NC n=6, *Fgl1^LWT^* WD n=8, *Fgl1^LKO^* WD n=9) and were compared by Two-way ANOVA followed by Holm-Šídák’s multiple comparisons test (A-D)or by Student’s t-test (D-M), *p<0.05, **p<0.01, ***p<0.001, ****p<0.0001.

**Figure 4:**
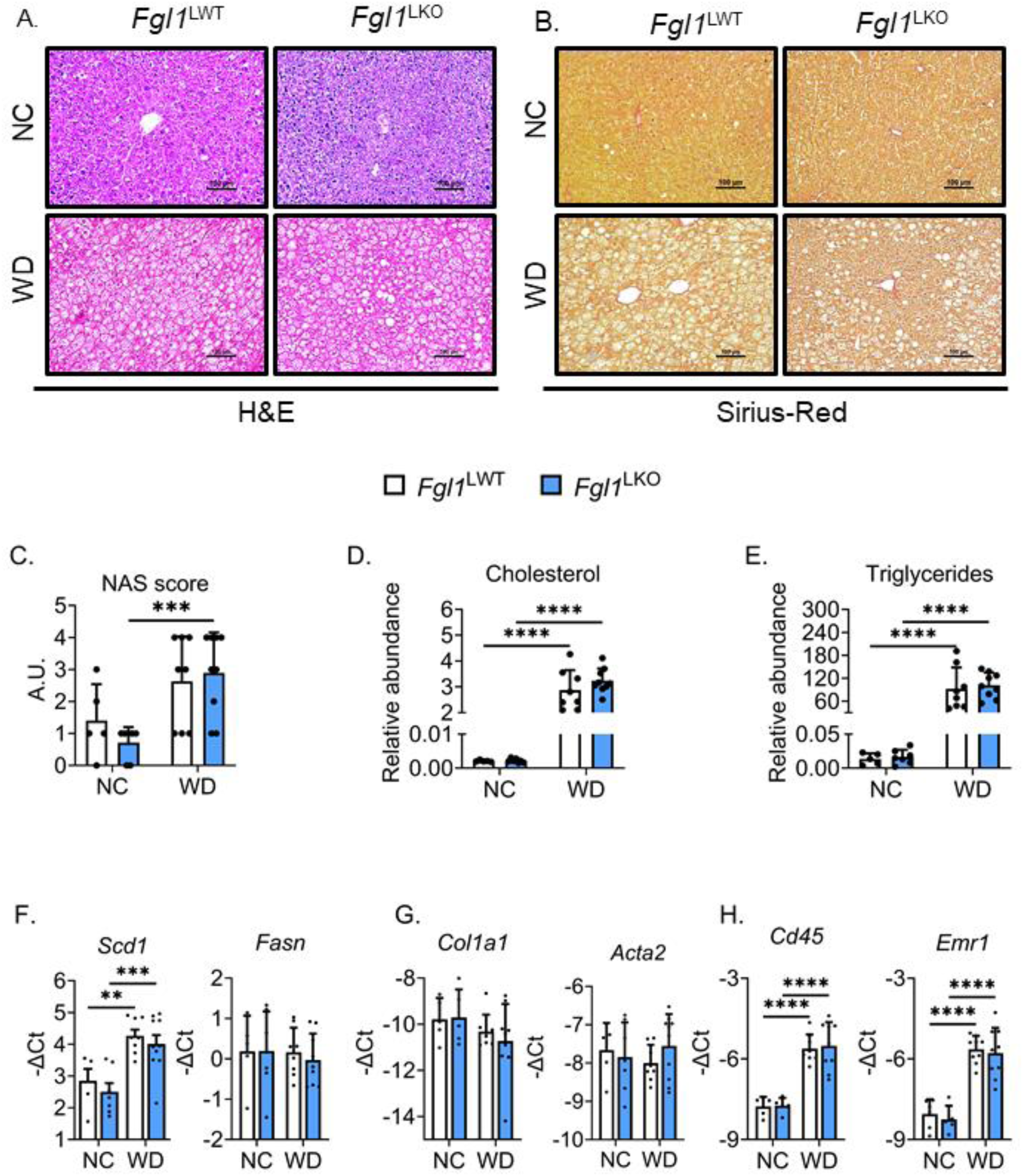
*Fgl1*^LWT^ and *Fgl1*^LKO^ mice displayed similar liver injury. Liver tissue sections were stained with hematoxylin and eosin (A) or Sirius red (B) and the Non-Alcoholic Fatty Liver Disease Activity Score (NAS, C) was computed. Scale bar is 100µm. Liver cholesterol (D) and triglycerides (E) content. mRNA expression of lipogenesis (*Scd1* and *Fasn*, F), fibrosis (*Col1a1, Acta2,* G) and immune cells markers (*Cd45* and *Emr1,* H) was measured by qPCR. Data are means ± SD (*Fgl1*^LWT^ NC n=5, *Fgl1*^LKO^ NC n=6, *Fgl1*^LWT^ WD n=8, *Fgl1*^LKO^ WD n=9) and were compared by Two-way ANOVA followed by Holm-Šídák’s multiple comparisons test, **p<0.01, ***p<0.001, ****p<0.0001.

### FGL1 deletion does not influence the progression of hepatocellular carcinoma

As FGL1 has also been described as a mediator of immune evasion (18), we assessed the effect of *Fgl1* deletion in the onset and progression of HCC using a protocol combining a western diet with weekly injection of CCl_4_ for 24 weeks (24). In this model both genotypes exhibited limited weight gain when fed a western diet (Figure 5A). After 24 weeks, both genotypes displayed similar liver to body weight ratio (Figure 5B) and circulating levels of ALT and AST (Figure 5C-D). However, we observed that 30% of *Fgl1*^LKO^ mice presented with macroscopic liver tumors (Figure 5E), suggesting an increased susceptibility to carcinogenesis in absence of FGL1. We next examined the extent of liver damages and fibrosis on histological sections stained with hematoxylin and eosin or sirius red and a NAS score was computed (Figure 5F-G). No difference was observed between *Fgl1*^WT^ and *Fgl1*^LKO^, indicating similar liver damages. mRNA levels of markers of inflammation (*Il6*), fibrosis (*Acta2* and *Col1a1)* (Figure 5H) and immune cells infiltration (*Cd45* for nucleated hematopoietic cells*; Ly6g* for monocytes, granulocytes and neutrophils*; Cd3e* for T-cells, Figure 5I) were similar between both genotypes. We found that *Emr1* (macrophages) expression was increased in *Fgl1*^LKO^ but the protein level was comparable when liver tissue sections were stained for F4/80 (Figure 5J). Thus, the deletion of *Fgl1* in hepatocytes does not influence the installation and progression of hepatocellular carcinoma.

**Figure 5:**
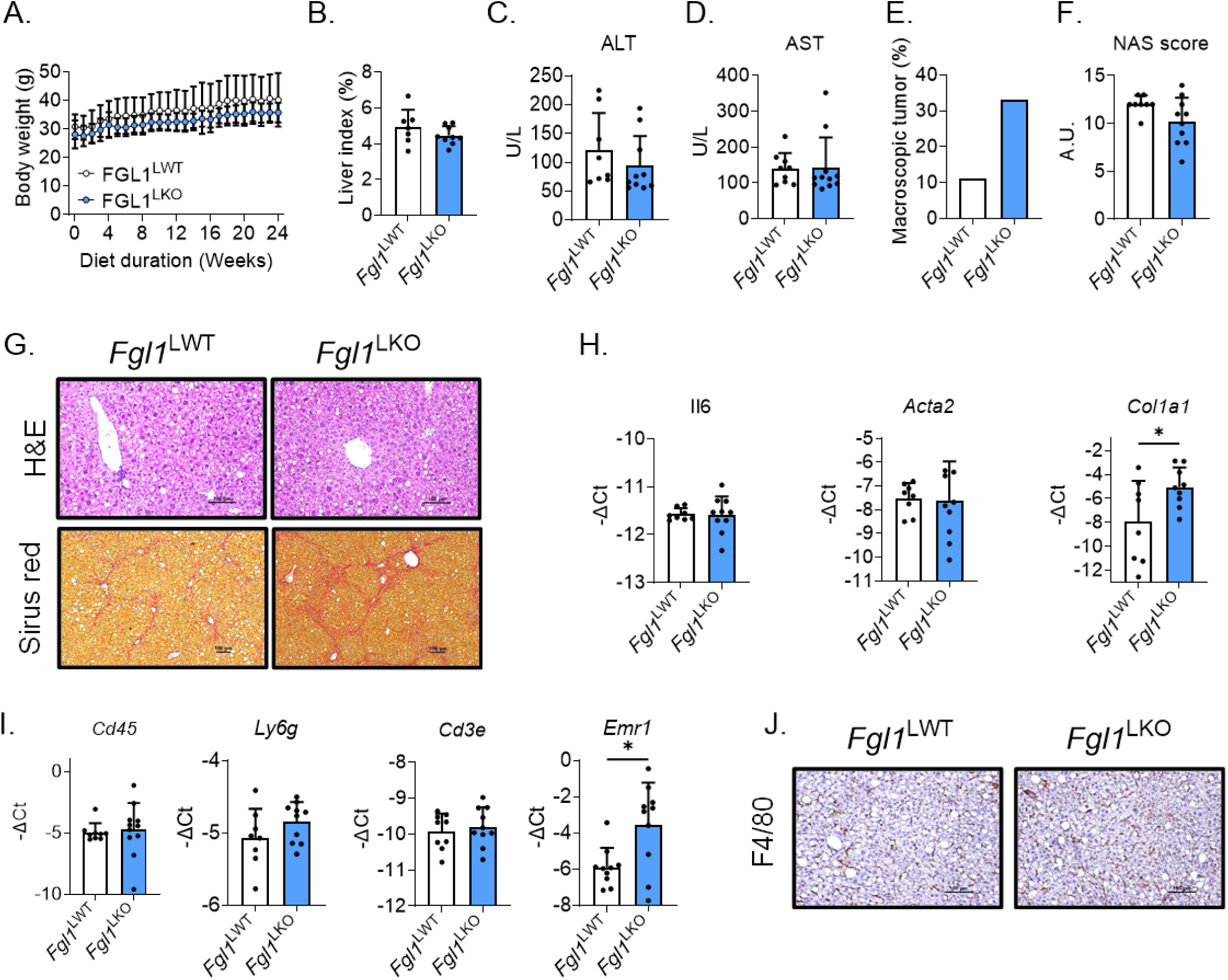
FGL1 does not contribute to the progression of fibrosis and hepatocellular carcinoma. *Fgl1*^LWT^ (n=11) and *Fgl1*^LKO^ (n=9) male mice were fed a western diet for 24 weeks and given weekly intraperitoneal injections of carbon tetrachloride (CCl_4_, 0.2µL/g). Body weight (A), liver weight relative to total body weight index (%) (B) and circulating alanine (ALT, C) and aspartate (AST, D) amino transferase levels. Percentage of observed liver macroscopic tumors (E). NAS score (F) and liver tissue sections stained with hematoxylin or picrosirius red (G). Liver mRNA expression of inflammation, fibrosis (H) and immune cells infiltration (I) markers were measured by qPCR. Liver sections were stained for F4/80 (J). Data are means ± SD and were compared by Two-way ANOVA followed by Holm-Šídák’s multiple comparisons test (A) or Student’s t-test (B-J), **p<0.01.

### *FGL1* expression is repressed during the progression of MASLD in mice and human

Retrospective analysis of published datasets from studies of MASH or HCC patients (GSE48452 (34), GSE164760 (35) and GSE14520 (36)) revealed that *FGL1* expression was significantly reduced in patients with MASH and HCC (Figure 6A-C). We therefore decided to study the time-course of *Fgl1* mRNA expression in the liver of mice fed a western diet for 8 weeks. We observed a gradual increase in steatosis and NAS score (Figure 6D-E) over 8 weeks but at this time-point circulating levels of ALT and AST were unchanged (Figure 6F). *Fgl1* mRNA expression was first induced after 4 weeks but decreased significantly 4 to 8 weeks after the mice were placed on western diet (Figure 6G). Interestingly, while the expression of the inflammatory marker *Socs3* and was unchanged (Figure 6H), *Fgl1* expression was inversely correlated with the expression of markers of ER stress (*Ddit3*) and immune cells infiltration (*Cd45*) (Figure 6I-J). These results indicate that *Fgl1* expression is repressed during the early phases of MASLD.

**Figure 6:**
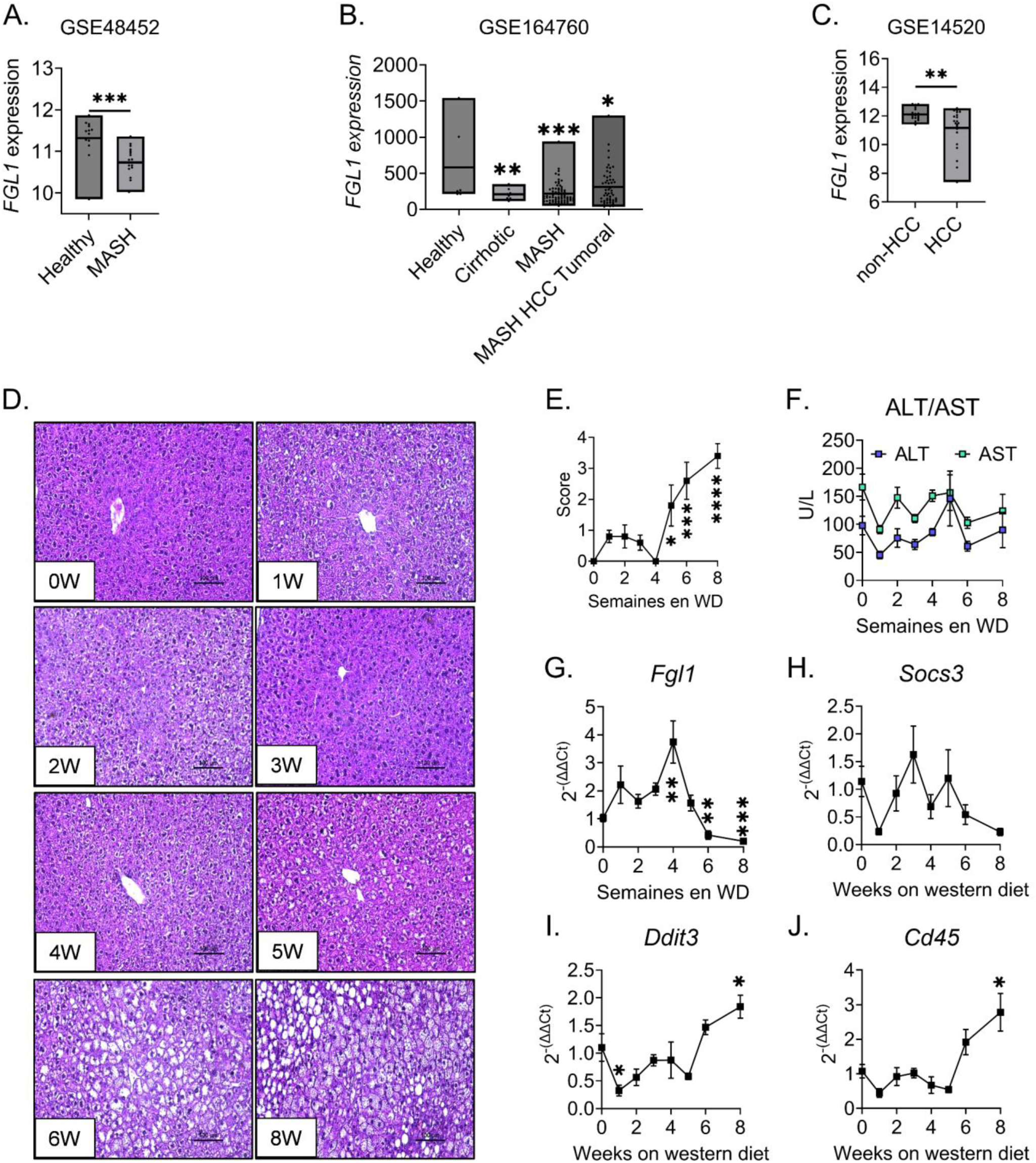
*Fgl1* expression is rapidly repressed during the onset of steatosis. Retrospective analysis of *FGL1* expression in published datasets GSE48452 (A), GSE164760 (B) and GSE14520 (C). Liver tissue sections stained with hematoxylin (n=5 per time-point) (D), NAS score (E) and circulating levels of ALT and AST (F) of mice fed a wester diet for 0-8 weeks. Corresponding liver *Fgl1* (G)*, Socs3* (H)*, Ddit3* (I) and *Cd45* (J) mRNA expression. Data are means ± SD were compared by Student’s t-test (A-C) or One-way ANOVA followed by Holm-Šídák’s multiple comparisons test (D-J), *p<0.05, **p<0.01, ***p<0.001, ****p<0.0001.

## DISCUSSION

Over the past decade, hepatokines have emerged as candidate contributing factors in the pathogenesis of MASLD (8), a complex pathology associated with multi-organ metabolic dysfunctions. Among those, the importance of FGL1 in the progression of MASLD remains speculative. While its official symbol is FGL1 for fibrinogen-like 1, it is also commonly referred to as hepassocin (HPS) or HFREP1 in studies that have yielded sometimes opposite conclusions depending on the strategy used to assess the potential function of FGL1 in metabolic diseases. These include transient knock-downs using shRNA, lentiviral-based overexpression, recombinant protein administration in combination with diet or genetically (*i.e.* Ob/Ob mice) - induced metabolic dysfunctions. Since FGL1 is secreted by hepatocytes, we examined for the first time the metabolic functions of hepatocyte-derived FGL1 using mice with specific ablation of *Fgl1* in hepatocytes. Similar to whole-body knock out (14, 15), *Fgl1*^LKO^ presented with increased adipose tissue deposits and slightly increase fasting glycemia compared to their littermate WT controls indicating that the metabolic functions of FGL1 are primarily attributable to hepatocyte-derived FGL1. As a non-significant trend towards reduced liver glycogen stores was observed, an increase in glycogenolytic activity may explain the difference in glycemia observed between genotypes. Suprisingly, only 7 genes that have never been previously linked to FGL1 were differentially expressed in the liver of mice lacking FGL1 compared to their WT controls. These genes do not encode for proteins related to FGL1 which excludes a potential compensation from homologous proteins. This clearly shows that FGL1 does not influence the liver transcriptional program. Although *Fgl1*^LKO^ and *Fgl1-/-* mice presented with a mild increase in fasting blood glycemia, the glucose tolerance is similar between WT mice and mice lacking FGL1 and the ablation of *Fgl1* had no effect on insulin signaling *in vivo.* Exogenous FGL1 administration seemed to act as an insulin sensitizer *in vitro* which contrasts with previous reports describing FGL1 as an inhibitor of insulin signaling (9, 11). Collectively, we show that FGL1 has little to no contribution in body metabolism in steady state conditions.

When fed a western diet for 16 weeks, a model that recapitulates the features of the pathology described in human (24), *Fgl1*^LKO^ mice exhibited slightly elevated ASAT, ALAT and triglyceride levels suggesting an increased sensitivity to diet-induced liver injury. However, we did not detect any difference in lipid accumulation in the liver and the MASH phenotype was comparable to their WT controls. In mice, administration of recombinant FGL1 or its overexpression using a lentiviral vector was previously shown to induce insulin resistance through the ERK1/2 signaling pathway whereas its lentiviral knockdown improved insulin resistance in mice fed a high fat diet or genetically obese Ob/Ob mice (9). The same group also used lentiviral vectors or short-hairpin RNA to show that, in mice fed a high fat diet for 12 weeks, overexpression of FGL1 increased hepatic lipid accumulation and NAFLD score whereas its knockdown reduced the steatosis and the NAS score (10). We observed a comparable increase in circulating transaminases levels in mice fed a western diet for 16 weeks compared to mice fed a normal chow. However, we reached opposite conclusions as the genetic deletion of *Fgl1* in hepatocytes was accompanied by an increase in ALT, AST and triglycerides levels compared to WT mice. Moreover, we did not observe any difference in ERK1/2 phosphorylation in the liver of *Fgl1*^WT^ and *Fgl1*^LKO^ and in primary mouse hepatocytes treated with FGL1. Our results are in line with a recent study showing that whole-body *Fgl1* knock out developed a more pronounced steatosis than their WT counterparts when fed a high fat diet for 16 weeks or a methionine-choline deficient (MCD) diet for 3 weeks (14). Daily injections of recombinant FGL1 (1mg/kg) for a week was sufficient to ameliorate the steatosis and insulin resistance in mice fed a high fat (16 weeks) or MCD (3 weeks) diet (14). Similarly, *Fgl1* silencing using siRNA enhanced liver injury induced by D-galactosamine in mice (31) whereas the administration of recombinant FGL1 to cynomolgus macaques limited the extent of liver injury induced by D-Galactosamine administration (37). Of note, we tested the antibodies used to assess liver FGL1 protein levels in mouse (9, 10, 14) and observed a similar protein profile in liver extracts or tissue sections from WT or *Fgl1-/-* mice (data not shown) indicating that the commercial antibodies to mouse FGL1 lack specificity.

Importantly, we found that *Fgl1* expression was significantly increased after 4 weeks on western diet (*i.e.* during the onset of the steatosis) and subsequently reduced after 6-8 weeks when the steatosis progressed. Two additional study described a similar bi-phasic regulation of *Fgl1* transcription during liver diseases. In mice fed a MCD diet, liver *Fgl1* mRNA expression was increased after a week and progressively decreased over 2-6 weeks of dietary challenge (14). A similar downregulation of *Fgl1* mRNA expression was observed in MASH patients and in the liver of 4-12-week-old *Ob/Ob* mice. FGL1 protein levels were presumably increased when mice were fed a high fat diet containing 45% lipids and significantly reduced after 24-40 weeks (14). An increase in FGL1 expression or protein levels was also observed in the liver of mice fed a high fat diet for 12 weeks (9, 10) but the mRNA expression data were not provided for these sets of mice. The transcription of *Fgl1* is regulated by inflammatory cytokines such as interleukin-6 in cooperation with the hepatocyte nuclear factor 1α (HNF1α) (14, 38) and the high-mobility group box 1 protein (HMGB1) which could explain the induction of *Fgl1* during the early phases (39) but we did not notice any change in the expression of the inflammatory marker *Socs3*. Similarly*, Fgl1* transcription is stimulated during ER stress (30) but the expression of ER stress target gene *Ddit3* inversely correlated with *Fgl1* expression. The present work support the most recent studies (14, 27, 40) suggesting that FGL1 may exert a protective function during the progression of steatosis but its exact contribution and the mechanism regulating *Fgl1* expression during metabolic alterations are still unknown.

Interestingly, 8-week-old *Fgl1*^LKO^ mice fed a normal chow displayed an increased adiposity compared to their WT controls. A similar phenotype was observed in *Fgl1-/-* mice by Demchev and colleagues (15). However, the same group that used lentiviral vectors to assess the contribution of FGL1 in MASLD described that lentiviral overexpression of *Fgl1* in epididymal fat pads increased the fat pads whereas its inhibition decreased the high fat diet induced adiposity (32). This further emphasizes the discrepancies between the genetic and lentiviral manipulation of FGL1 suggesting that the absence of FGL1 during development and at birth could predispose to a different response during metabolic cues in comparison to a transient inactivation. As the expression of *Fgl1* in adipose tissue is very low (41), the phenotype could also be attributable to lentiviral vectors off-target effects. The molecular mechanisms involved in the inter-organ dialogue between hepatocyte-derived FGL1 and the adipose tissue therefore remain uncertain.

FGL1 has also been described as a mediator of immune evasion in certain cancers such as non-small cells lung cancer or metastatic melanoma (18). The binding of FGL1 to its receptor lymphocytes activating gene 3 (LAG3) at the cell surface of activated T lymphocytes promotes T cells exhaustion and inactivation and prevents their anti-tumoral activity (18). While we observed a trend toward more solid tumors in *Fgl1*^LKO^ mice compared to *Fgl1*^LWT^ mice, both genotypes exhibited a similar HCC phenotype. As the expression of LAG3 is low in the liver (42), the FGL1/LAG3 interaction may not contribute to liver cancer. Moreover, this mechanism has been recently questioned (19, 20) and FGL1 may only have a weak tumorigenic activity.

FGL1 has been originally described as an hepatocytes mitogen (12, 31, 43, 44). Although knockdown of FGL1 during zebrafish embryogenesis using morpholino reduced the proliferation of hepatocytes and the size of the liver, no change in liver weight was found in *Fgl1-/-* mice. In human, the downregulation of FGL1 is observed in 20% (18) to 60% (40) of liver cancer and its expression correlated with the degree of tumor differentiation. FGL1 is located on chromosome 8p, a region where cumulative allelic loss have been positively correlated with aggressive tumors (17, 45). Conversely, high *FGL1* expression was significantly associated with larger tumor size and liver cirrhosis (46). The study of FGL1 protein level in human will await the development of new analytical tools and standardized methods. Indeed, published results of FGL1 measurement using commercial ELISAs should be interpreted with caution as the reported baseline levels in healthy patients varied by several order of magnitude.

## Data availability statement

The data that support the findings of this study are available in the Materials and Methods and Results section of this article.

## Conflict of interest

The authors declare that they have no conflicts of interest with the contents of this article.

## Authorship Contributions

JP, LC, CMPM, PP, AD, MS, JDS performed experiments and analyzed data. JP, HG and LK designed and supervised the study, analyzed data and wrote the manuscript. All authors edited the manuscript.

## Acknowledgments

The authors thank Céline Deraison (Inserm U1220) and Nicolas Blanchard (Inserm U1291) for providing mice expressing the flippase 1 (FLP1), members of the Inserm US006 animal, histopathology and phenotyping facility facilities (Toulouse) and the platform Aninfimip, an EquipEx (‘Equipement d’Excellence’) supported by the French government through the Investments for the Future program, Nancy Geoffre and Justine Bertrand-Michel from MetaToul (Toulouse metabolomics & fluxomics facilities, www.metatoul.fr) which is part of the French National Infrastructure for Metabolomics and Fluxomics MetaboHUB-ANR-11-INBS-0010 (doi.org/10.15454/VRJK-KY76). Support for this work was provided by the French National Research Agency (ANR-16-ACHN-0002-01 and), the region Occitanie and by the European Research Council (ERC) under the European Union’s Horizon 2020 research and innovation program (grant agreement no. 715491) to LK and by ARC (ARCPOST-DOC2022020004813) and FRM (ARF202209015711) postdoctoral fellowship to JP.

## Notes

### Competing Interest Statement

The authors have declared no competing interest.

### Summary of Updates

We are now providing an extensively revised manuscript that includes a new figure and several additional panels showing: - a comparison of Fgl1 mRNA expression in hepatocytes vs adipose tissue - a comparison of the transcriptomic profiles of livers from Fgl1LWT and Fgl1LKO mice by RNA-sequencing - expression data of marker genes of gluconeogenesis (Pck1, G6pc) in the liver and of lipolysis (Lipe, Pnpla2), circulating triglycerides hydrolysis (Lpl) and fatty acid transport (Cd36) in epidydimal and inguinal adipose tissue of Fgl1LWT and Fgl1LKO mice - blood glucose and ketone in Fgl1LWT and Fgl1LKO mice - liver glycogen levels in Fgl1LWT and Fgl1LKO mice - an assessment of insulin signaling in the liver of Fgl1LWT and Fgl1LKO mice and in hepatocytes treated with FGL1 by western blot (AKT and ERK1/2).

